# Erythrocyte ghosts and Plasma beads as Antigen peptide delivery systems in animals and humans

**DOI:** 10.1101/2023.03.09.531854

**Authors:** M. Rizwan Jameel, Hina Jameel, Saad Mustafa, Mohd Faisal Khan, Mohammad Misbah

## Abstract

Erythrocytes are the most plentiful cells of the human body with desirable morphologic and physiologic characteristics and their function in drug delivery has been exploited comprehensively. These cellular carriers, containing significant biodegradability, biocompatibility, and duration in circulation, can be burdened by a wide spectrum of compounds of therapeutic interest Carrier erythrocytes holding drugs, enzymes or peptides, Cas, and Nanoparticles can be used as a delivery system that approval to changes in the kinetic behavior and selective bio-distribution of the substances encapsulated. Carrier erythrocytes have been exploited for numerous potential applications, including the intravenous slow release of drug targeting, therapeutic agents and enzyme therapy to a reticuloendothelial system (RES). Hypotonic dialysis is the process most usually utilized in the formation of erythrocytes of carrier, however many features concern the give up and the erythrocytes ghost characteristics obtained with this technique. Attempt was also completed observe but erythrocyte regular release entrapped substances/antigens with additional entrapment in the plasma beads, improves the protein immunogenicity. The glycoenzyme invertase of large molecular weight was utilized a model substance and entrapment of invertase consummate with the dialysis method. Enzyme entrapment was examined in derived erythrocytes from rabbits, goats and human. Intended for the immunological studies were used in the rabbits. Invertase entrapped erythrocytes rabbit, where point out was additional encapsulated in the plasma beads and a contrast of immunogenicity of lose invertase, entrapped invertase in erythrocytes and that entrapped in RBCs and additional in encapsulated the plasma beads were examined. Antibodies production was investigated are different periods such as 12, 20, 20 and 35 days.

## INTRODUCTION

For the earliest instance, erythrocytes were expressed in samples of human blood through the Dutch scientist Lee Van Hock in 1674. Regarding hundred years afterward, Howson establish that these cells are plane discs rather than similar to the globules explained by Lee Van Hock, and early in 19h century [1], Hope Seyler recognized hemoglobin in erythrocytes and its vital function in oxygen release to different tissues. In the human body, the mature forms of erythrocytes are normally biconcave disks non-nucleated with a middle pallor. The biconcave form of erythrocytes gives a large ratio of surface-to-volume for the delivery of oxygen and enhanced flexibility within movement during narrow capillaries. The erythrocyte’s average content in women and men is 4.8* 10^6^ and 5.4* 10^6^ per ml, respectively [2]. Erythrocytes comprise potential highly biocompatible vectors used for the delivery of various bioactive substances, as well as proteins and drugs. The first effort at the encapsulation of drugs within erythrocyte cells extends to the peptides and enzymes through therapeutic activity [3–5].The utilization of red cells as transporters of drugs constitutes a field of effort that has hardly been investigated, particularly when evaluated to the carrier structures. Many procedures are available for the entrapment of pharmaceuticals in erythrocytes. These include procedures based on reversible perturbation of the membrane by physical or chemical strategies including dielectrics breakdown, osmotic hemolysis, etc [6–8]. Irrespective of the process used, the most select characteristics for successful compound entrapment require the drug to contain a significant amount of water solubility, resistance alongside degradation inside erythrocytes, lack of chemical or physical interaction with the membrane of erythrocyte, and fine distinct pharmacodynamic and pharmacokinetic properties. Hypotonic hemolysis was primarily investigated for the chemicals encapsulation into erythrocytes and is the fastest and simplest. In this technique, an amount of packed erythrocytes is diluted through from 2 to 20 volumes of aqueous resolution of a drug. The tonicity of the solution is after that restored through the addition of a hypertonic buffer. The mixture result is after that centrifuging, discarding the supernatant and washing the pellet with the buffer of isotonic solution [7]. The chief disadvantages of this technique contain low entrapment effectiveness and a significant loss of hemoglobin and extra components of the cell. This decreases the half-life of the circulation of the loaded cells. RES macrophages are phagocytosed readily and thus may be utilized to target organs RES. The Diluted hypotonic is utilized to load enzymes for instance glucosidase, galactosidase, arginase and asparaginase and bronchodilators for example salbutamol [6, 8].

A network fibrin is a scaffold the first cell comes across while it carries outs its function within curative trauma wounds or normal tissue of other insults. Bind platelets and pull on the filament of the fibrin when they collect in a network [9] macrophages and neutrophils [10] connect to fibrin as they abode to injury sites disposing of infectious agents and dead tissues, they have violated the barrier epidermal, and fibroblasts initial secure to fibrin like they go in the lesion site to restructure the tissue [11, 12]. Distinct the basement membranes and extracellular matrices shaped by laminin, proteoglycans and collagen, which collect gradually in a prepared way dictated through the cells to secrete them, a rich ingredient of plasma of blood, fibrin gels accumulate quickly through an adapted reaction of polycondensation [13] from fibrinogen, immediately the Thrombin protease is turned on in the cascade clotting and starts to eliminate the fibrinogen polypeptides of the part that stop its polymerization spontaneous. The fibres branching 3 dimensional networks is shown in figure 1, that hydrogels form with bulky elastic moduli at very low volume of polymer fraction. Fibrin gels produced from purified isolated plasma proteins obtain mechanical and structures properties that are very comparable to the blood clot. Additionally, the clot of fibrin is biodegradable, being impartial in a systematic with the fibrinolytic system way [14]. The cells presence, particularly the ones for instance platelets that encourage inner strains in network of the fibrin [15], changes blood clots of the mechanical properties in vivo, but the fibrin strands structures and the networks geometry, they structure are comparable to those produced in vitro. While platelets do as well concern structure of fibrin since their membrane have fibrin (fibrogen) required enzyme complexes and integrins to facilitate thrombin activate, adding of isolated cells. To provide wound repair, it is frequently attractive to nearby transport specific tissue growth of factors in a manner of controlled. Fibrin is an engaging delivery vehicle drug since it can be inserted wherever it gels into situ; it is naturally ruined and excited the body’s own healing injury response. This is an additional function wherever fibrin has established practical. Suspended keratinocytes in fibrin were effectual in full thickness reconstituting injures in both subjects of mouse and human, although skin coated fibroblast, microbeads of fibrin reduced the time for tissue formation of granulation [16, 17].

**Figure 1.**
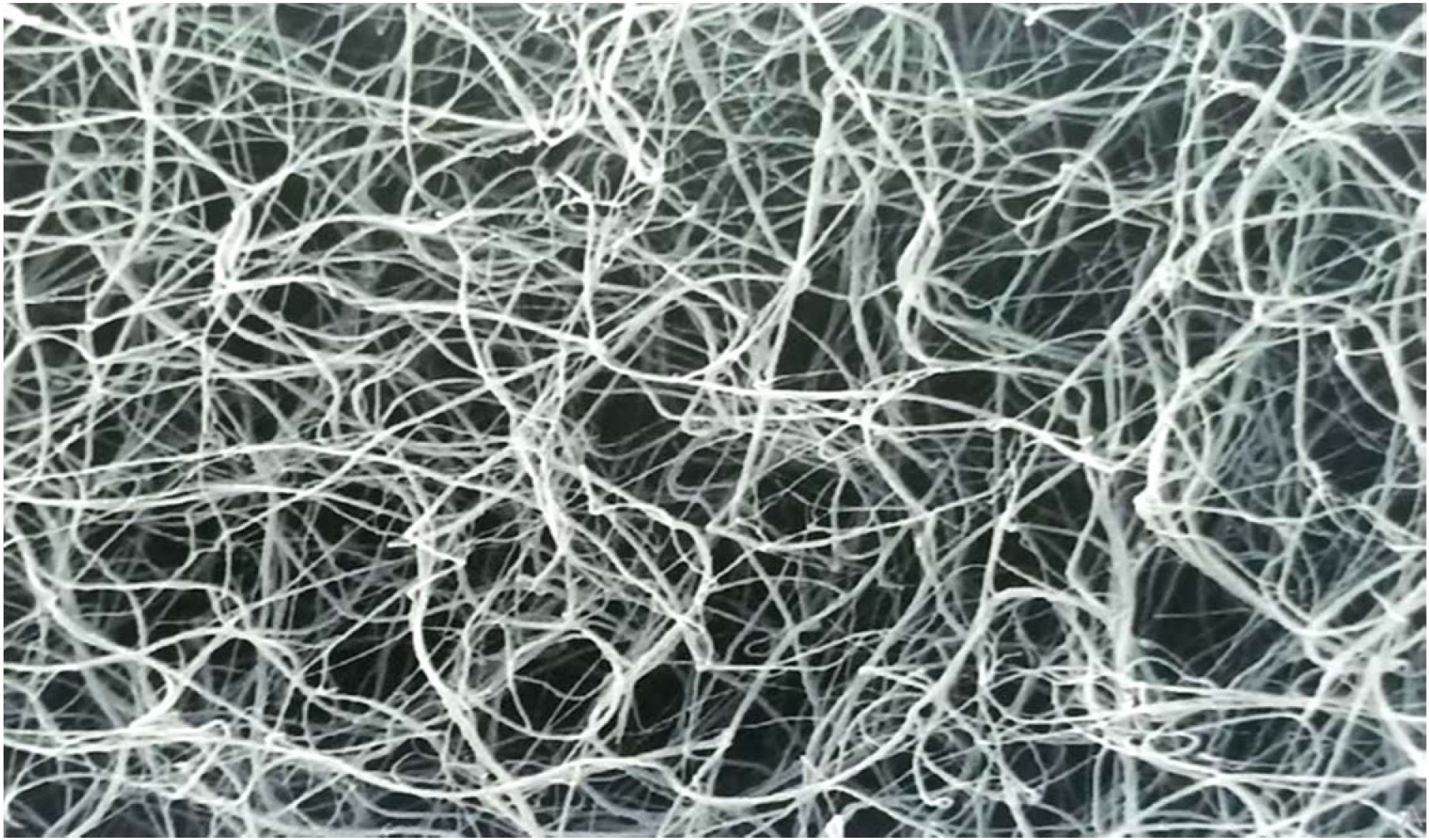
Scanning electron micrograph human fibrinogen polymerized with thrombin at pH 7.4 and 150 mM NaCl.

## MATERIALS AND METHODS

### Estimation plasma protein

The procedure described by Lowry et al (1951) [18] depends on the exchange of Cu^2+^ to Cu in alkaline conditions. The reactions product in a well-built color blue, which bases partly on the tryptophan and tyrosine content. To sample/standard 0.1 mL was added to distilled water 0.9 mL and adds 5 mL of fresh copper reagents. The solutions were permitted to situate at room temperature with Folin’s reagent (0.1 ml) for 10 minutes with a vortex mixer, and permit the mixture to situate at room temperature for 30-60 minutes. The absorbance was read at 550 nm. An absorbance of the standard curve of like a purpose of the original concentration of protein prepared was used to conclude the unknown protein concentrations.

### SDS PAGE for Plasma Protein and Erythrocytes membrane

Analysis of SDS-PAGE was performed as illustrated by Laemmli (1970) [19] in a mini-gel electrophoresis vertical apparatus (ATTO, Japan) with 10% (w/v) proteins of resolving gel, 8% (w/v) erythrocytes membrane of resolving gel and 5% (w/v) stacking gel of acrylamide. Electrophoresis was accepted, to begin with at 9 mA to make possible the sample stacking and existing was afterward increased to 18 mA. Electrophoresis was performed in anticipation of the tracking dye attained at the base of the gel. The gels were subsequently removed from the plates of gels and stained with Coomassie Brilliant Blue R-250 0.1% (w/v) (equipped in methanol, water and glacial acetic acid in 10:45:10 ratio respectively). The gels were destained by a solution including of each ten parts methanol and glacial acetic acid and eighty parts of water.

### Invertase encapsulation into erythrocytes by hypotonic dialysis method

5 ml of packed erythrocytes which had been washed by centrifugation at 500 Xg for ten minutes thrice in 154 mM NaCl, the buffer of 10m M Tris (pH 7.4). The cells were added to a dialysis bag along with invertase and dialyzed at a 60% hematocrit against 1 liter of purified distilled water for two hours. The NaCl concentration was subsequently raised to 154 mM by the addition of adequate concentrated NaCl solution to facilitate resealing and the resealed ghosts were permitted to stand for at least 30 minutes at room temperature and then washed thoroughly to remove unentrapped material

### Preparation of plasma beads

Fresh was blood collected from rabbits in the presence of 2.7% (w/v) EDTA solution (in a volume ratio 9: l), EDTA in normal saline to prevent blood clotting. Blood was centrifuged at 500 Xg for 10 minutes and the supernatant plasma was collected and kept at −20 °C till use. 250 ul plasma was mixed with 50 ul of the substance to be trapped and 22 ul CaCl, (500 mM). 5.0 ul aliquots of the mixture were placed as a droplet more than covered a glass slide with Parafilm. The glass slides were then stored in a moist chamber. The moist chamber was a large-sized Petri plate, with wet sterilized filter sheets on both the inner surfaces. Further wet cotton was also located in the lower plate. Now, the moist chamber with beads was placed in an incubator at 37°C. After the polymerization of the fibrin network, the biomolecule entrapped into the beads is picked up with surgical blades and placed in a normal saline solution. With two items of washing in normal saline and two items of washing in phosphate-buffered saline (PBS)-pH 7.4, they were finally suspended in phosphate buffer (20 mM, pH 74)

### Invertase assay

The assay method explained by Bernfield 1955 [20] was followed to assay invertase. The mixture of the assay was in a total volume of 300 ul such as 150 ul of 0.2 M buffer of sodium acetate (pH 4.9), 100Ļul of the buffer having invertase and 50 ul of 0.5 M sucrose. The samples were kept warm for 10 minutes at 37°C and the reaction stopped through the count of 20 μl of 0.5 M sodium phosphate (pH 7.0). The mixture assay was kept back in a boiling-temperature water bath for 5 minutes. Later than cooling, 1,0 ml of 1 % (w/v) di-nitrosalicylic acid reagent (DNS) was extra and the tubes were reserved at room temperature for 5 minutes followed by 5 minutes of incubation at 100 °C temperature in a water bath. Three (3) milliliters of distilled water were added into each tube in order to make up the volume to 5.0 ml and the color was examined at 540 nm. Invertase for one unit is the quantity that converts 1.0 μmole of sucrose to fructose and glucose per minute at 37 °C.

### Enzyme Linked Immunosorbant Assay (ELISA)

An Antibody raised adjacent to invertase was observed with ELISA. Three animals with a usual weight of 1.7 kg were used in three separate groups. One group received free invertase in buffer subcutaneously and other groups received erythrocyte-encapsulated invertase and invertase encapsulated into erythrocytes encapsulated into plasma beads. The dose of the antigen was 600 pgs in each case. A vaccination dose of 200 ugs was given during the same routes 35 days after primary immunization. Collected sera at intervals regular and ELISA executed to study antibody response in various animals. The microtitre plates were coated with antigen (5 μgs of invertase /100 μl of coating buffer) in a microtitre plate and kept overnight at 4°C. The coating buffer comprised 0.05 M sodium bicarbonate-carbonate buffer (pH 9.4). The plates were washed thrice in phosphate-buffered saline (pH 7.0) having 0.05% Tween20 (PBST) and blocked with 5% (w/v) milk skimmed in PBS for 2 h at room temperature. The washed plates were incubated with 100ul/well sera diluted in blocking buffer for 2 hours at 37°C, washed thrice with PBST. The secondary antibody diluted 1:1000 in the blocking buffer was then incubated for 40 minutes. The substrate solution was prepared by adding 10 μl of 30% H_2_O_2_, and 10 mg O-phenyl diamine to 12 ml 0.1 M sodium citrate-citric acid buffer, pH 4.6. 100 μl of the substrate solution was added to each well of the microtitre plate after secondary antibody washings. The reaction was stopped by adding 50 μl of I M H_2_SO_4_, Absorbance was deliberate at 450 nm.

### Isolation of erythrocyte ghost membranes

Prepared Ghosts were with the methods of Steck et al. 1970 [21], which followed the principle of hypotonic lysis defined by Dodge et al. 1963 [22]. Phase contrast microscopy was used to observe the procedure. Newly drawn blood was mixed with 2.7% (w/v) EDTA with are a ratio of 9:1, then diluted with an equal volume of cold 5 mM sodium phosphate [pH 8 (5P (8)]-0.15 M NaCl. The cells were washed thrice washing by centrifuging for 2000 rpm at 10 minutes and were lysed by mixing quickly 1 ml portion into 20 ml of cold 5P (8). The suspensions were centrifuged for 10,000 at 30 minutes and were thrice washed and collected by centrifugation and the ghost membrane button. The membrane was subjected to SDS-PAGE for analysis of polypeptide composition [19].

## RESULT

### 1. Entrapment invertase

#### 1.1. Rabbit erythrocytes

Knowledge of the optimum conditions essential for the loading of enzymes is important before the enzyme-loaded rabbit erythrocytes can be used as an antigen. With this objective, we have studied the effect of added invertase concentration on enzyme loading in the erythrocytes as well as on entrapment efficiency and cell recovery. There is a consistent increase in the amount of entrapped invertase with an increase in the concentration of added enzyme over the range investigated. Further increase in the amount of enzyme resulted in no significant further increase of enzyme entrapment in rabbit erythrocytes. The data on this entrapment of the invertase in rabbit erythrocytes are exposed in Table I and Figure 2. These were concentration-dependent enhancements in the amount of invertase entrapped in the erythrocytes initially which gradually slopped off and beyond 600 U/0.1 ml, the entrapment value did not rise significantly. As is also evident from the data entrapment efficiency decreased gradually with an increase in added invertase.

**Figure 2:**
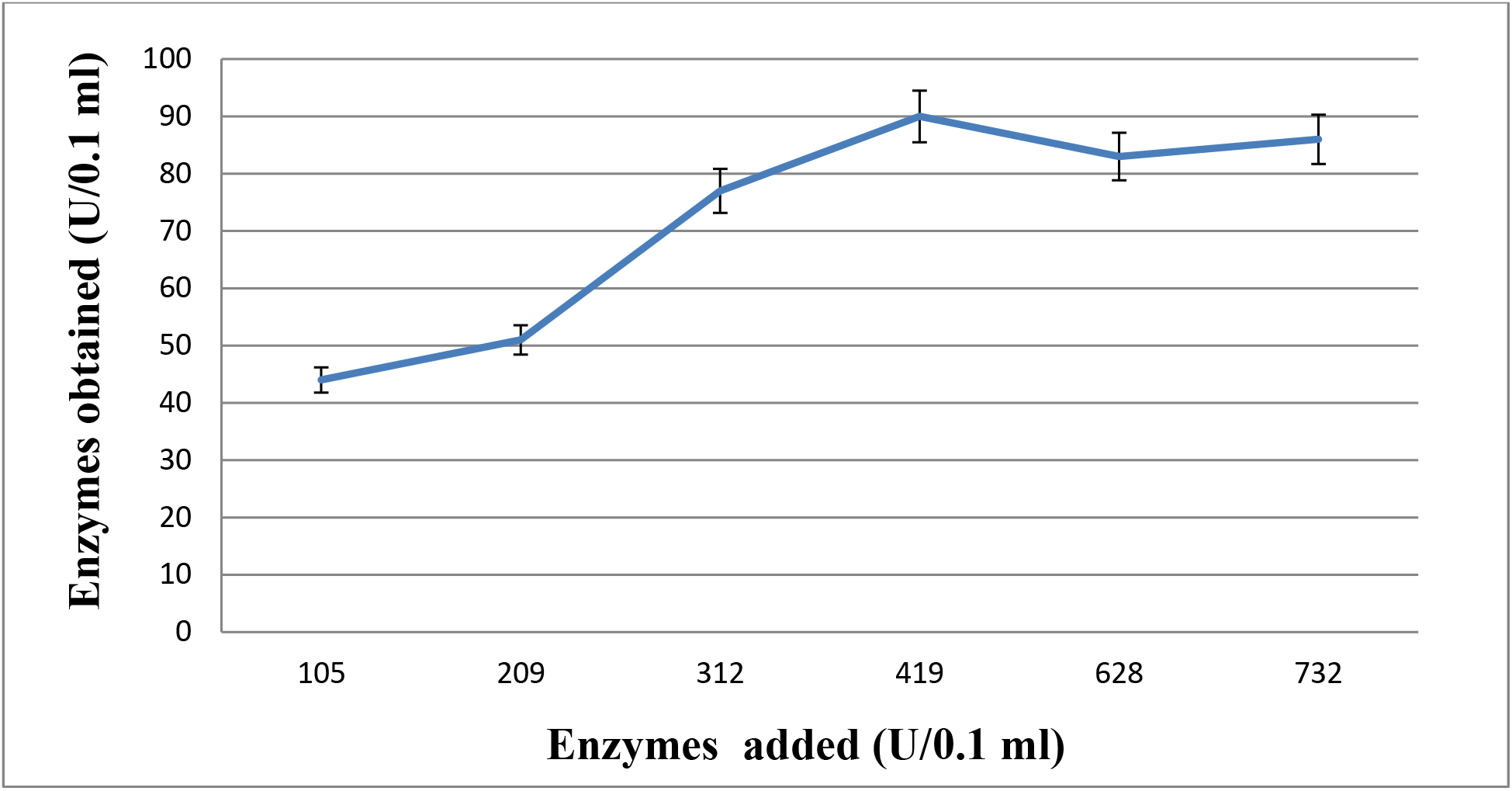
Effect of concentration of added invertase an enzymes in rabbit

**Table 1.**
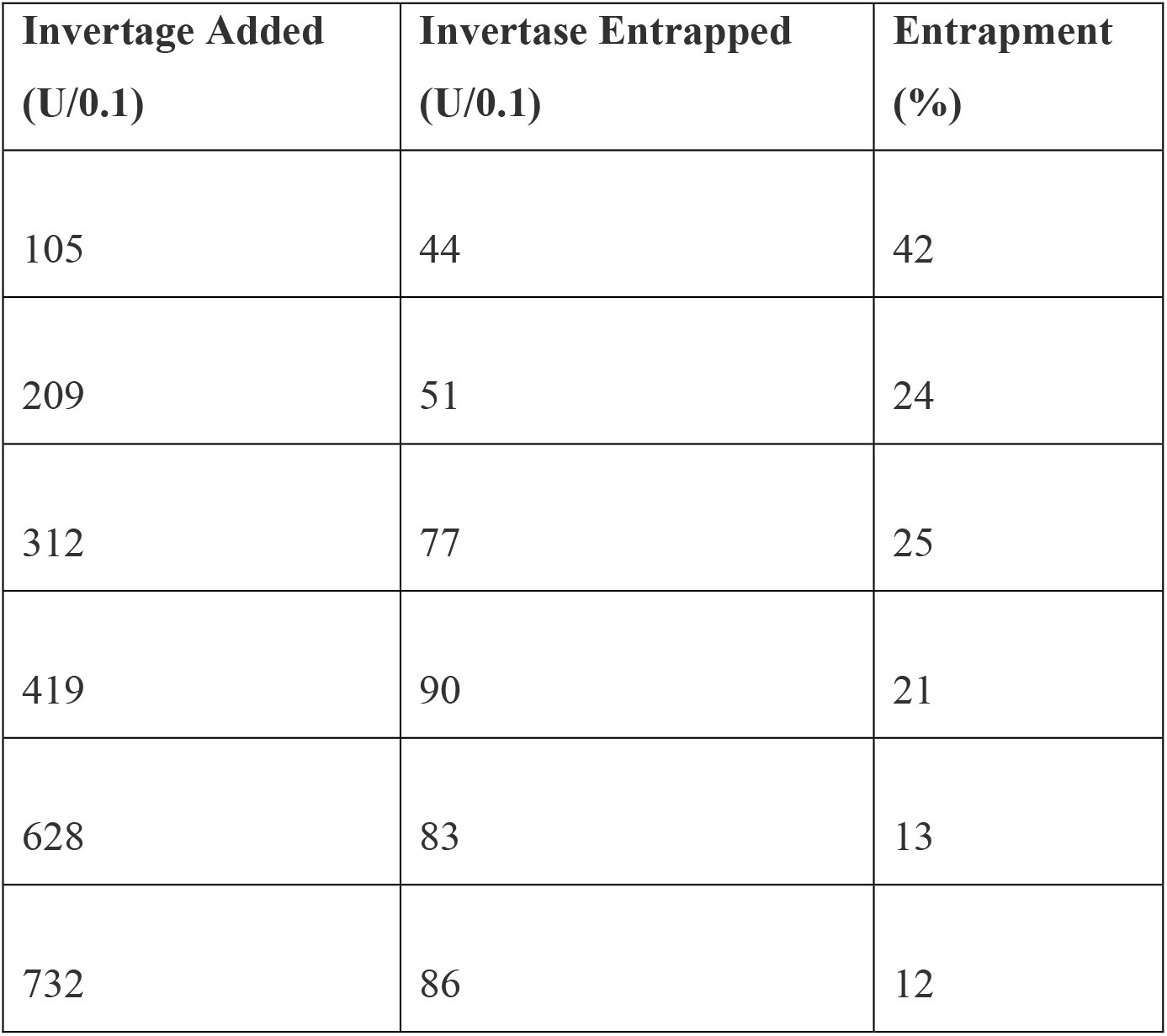
Invertase entrapment in rabbit erythrocytes.

The red blood cells were washed and enzyme entrapment performed using 0.7 ml of cell as described in methods each value is the average of atleast 2 determination.

#### 1.2. Goat erythrocytes

As from figure 3, there is a constant enhancement in the quantity of entrapped invertase by enhancing the concentration of additional enzymes up to 372 U/0.1 ml. The entrapment efficiency shows a slight decrease at each step up till the optima but that can be compensated for by the increase in final loaded packed cells in terms of U/0.Iml as can be seen in table 2.

**Figure 3:**
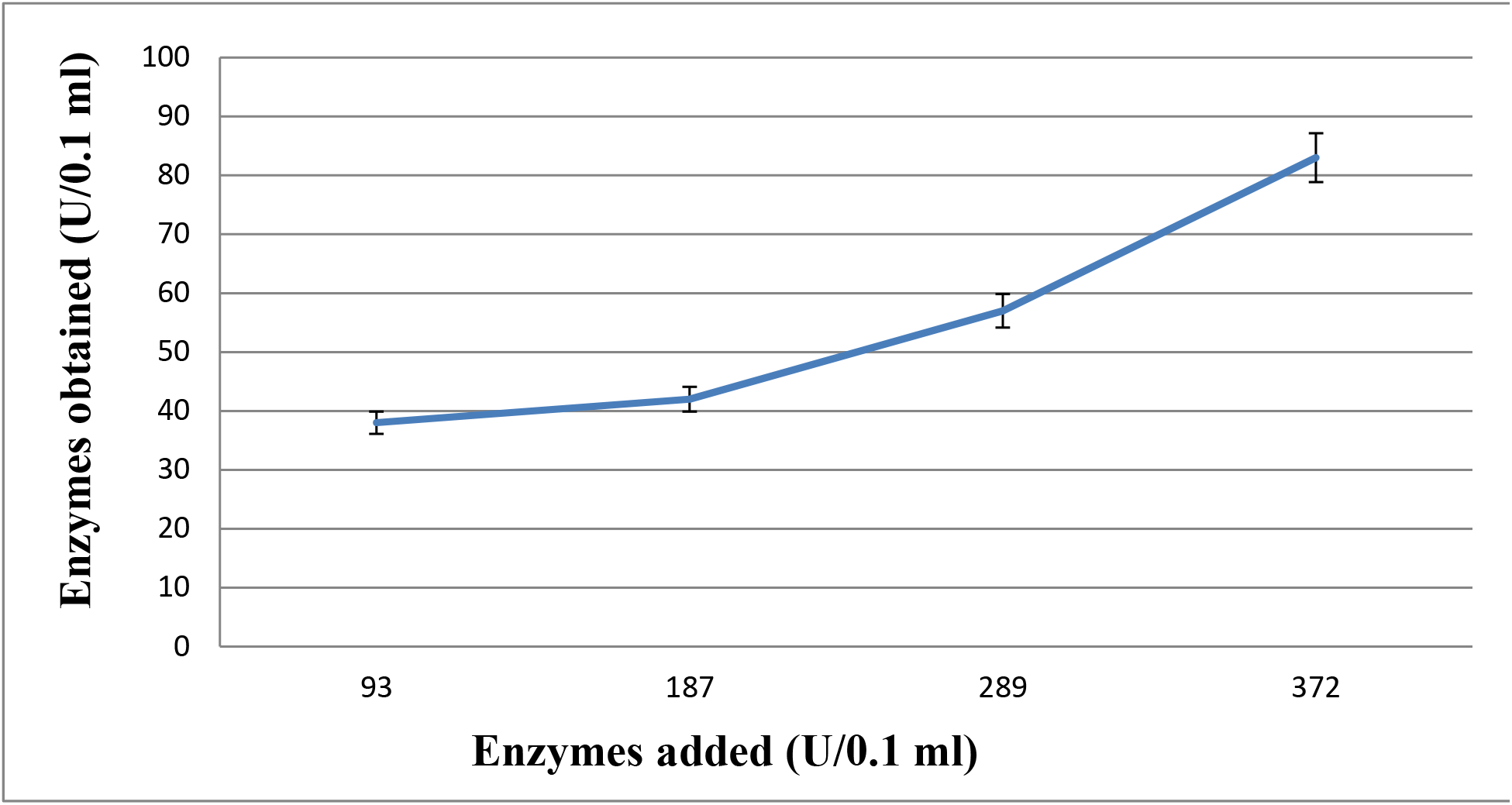
Effect of concentration of added invertase an enzymes in goats

**Table 2.** Invertase entrapment in goat erythrocytes.

| **Invertase Added**<br>**(U/0.1 ml)** | **Invertase**<br>**Entrepped (U/0.1 ml)** | **Entrapment**<br>**(%)** |
| --- | --- | --- |
| 93 | 38 | 41 |
| 187 | 42 | 22 |
| 289 | 57 | 20 |
| 372 | 83 | 22 |

The red blood cells were washed and enzyme entrapment performed using 0.7 ml of cell as described in methods each value is the average of atleast 2 determination

#### 1.3. Human erythrocytes

The data on this entrapment of invertase in human erythrocytes is shown in Table 3 and Figure 4. This was concentration dependent enhance in the quantity of invertase entrapped in the erythrocytes initially which gradually slopped off and beyond 235 U/O.1 ml, the entrapment value did not rise significantly As is also evident from the data entrapment efficiency decreased gradually as with increase

**Table 3.** Invertase entrapment in human erythrocytes.

| **Invertase Added**<br>**(U/0.1ml)** | **Invertase Entrapped**<br>**(U/0.1 ml)** | **Entrapment**<br>**(%)** |
| --- | --- | --- |
| 79 | 27 | 34 |
| 157 | 51 | 32 |
| 235 | 55 | 23 |
| 314 | 38 | 12 |

**Figure 4:**
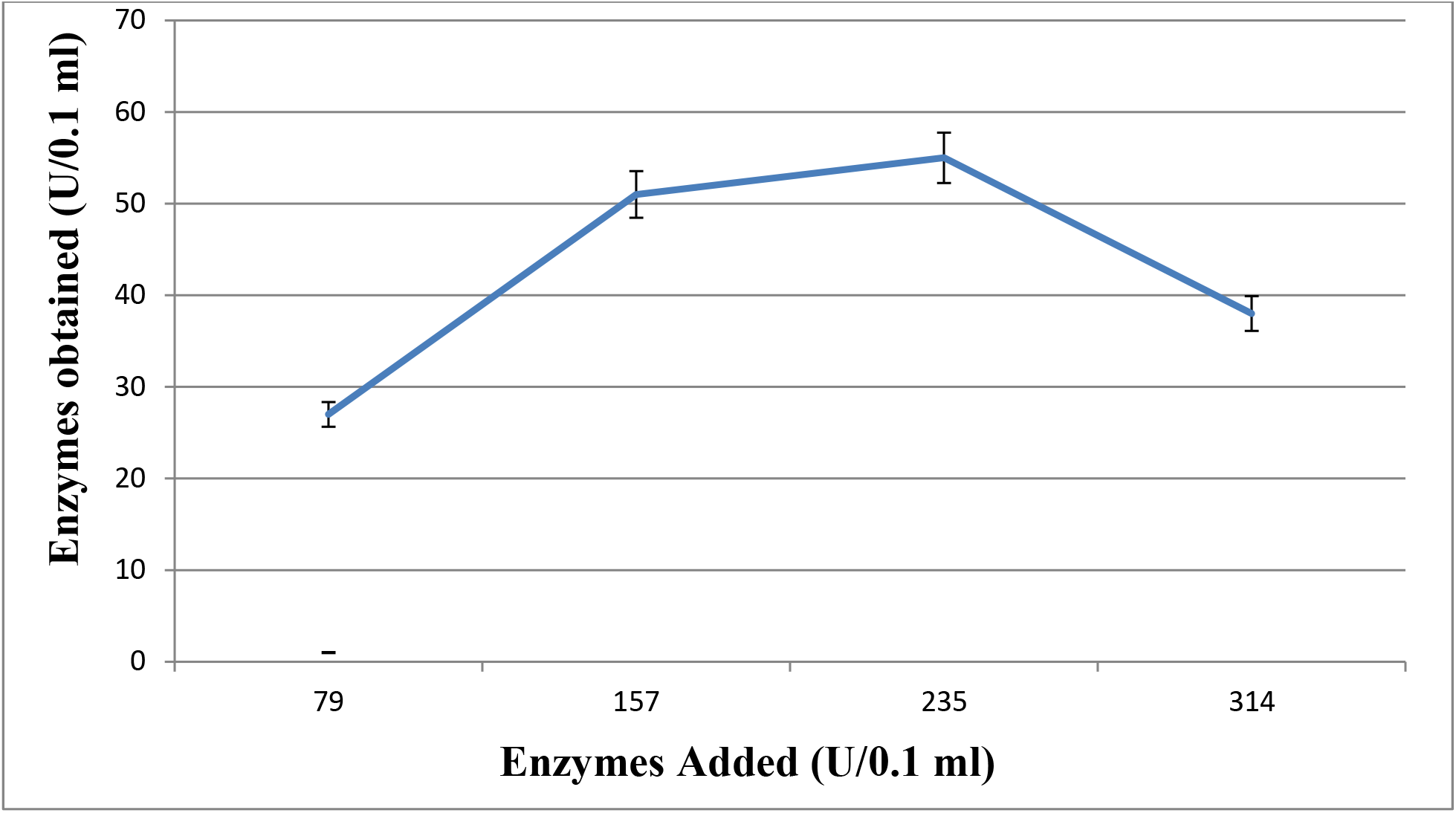
Effect of concentration of added invertase an enzymes in human erythrocytes.

The red blood cells were washed and enzyme entrapment performed using 0.7 ml of cell as described in methods each value is the average of atleast 2 determination.

## 2.0. Preparation of erythrocyte ghost and entrapment of enzyme loaded erythrocytes in the plasma beads

Figure 2 shows the polyacrylamide gel electrophoresis of the rabbit and human erythrocyte membranes. The membranes were characteristic of both large and small molecular weight proteins. One can see bands corresponding to Spectrin (94 KD), Band 3 (64 KD) and Actin (41 KD) and other beads were determined using the calibration curve prepared using the standard markers. The preparation of plasma beads and a procedure developed in this laboratory [23]. As described in the methods the procedure involves mixing of plasma attained for collected blood in presence of EDTA, mixing with the antigen-loaded erythrocytes and inducing clothing by addition of CaCl2. In order to prepare more stable beads, the plasma was concentrated before the addition of CaCl2. As exposed in figure 6 adding of the growing quantity of Sephadex G-10 the plasma effect in the concentration of the same. The protein concentration increased from 162 to 185 mg/ml when 50 mg Sephadex −10 was added to 250 μl plasma (P).

**Figure 5:**
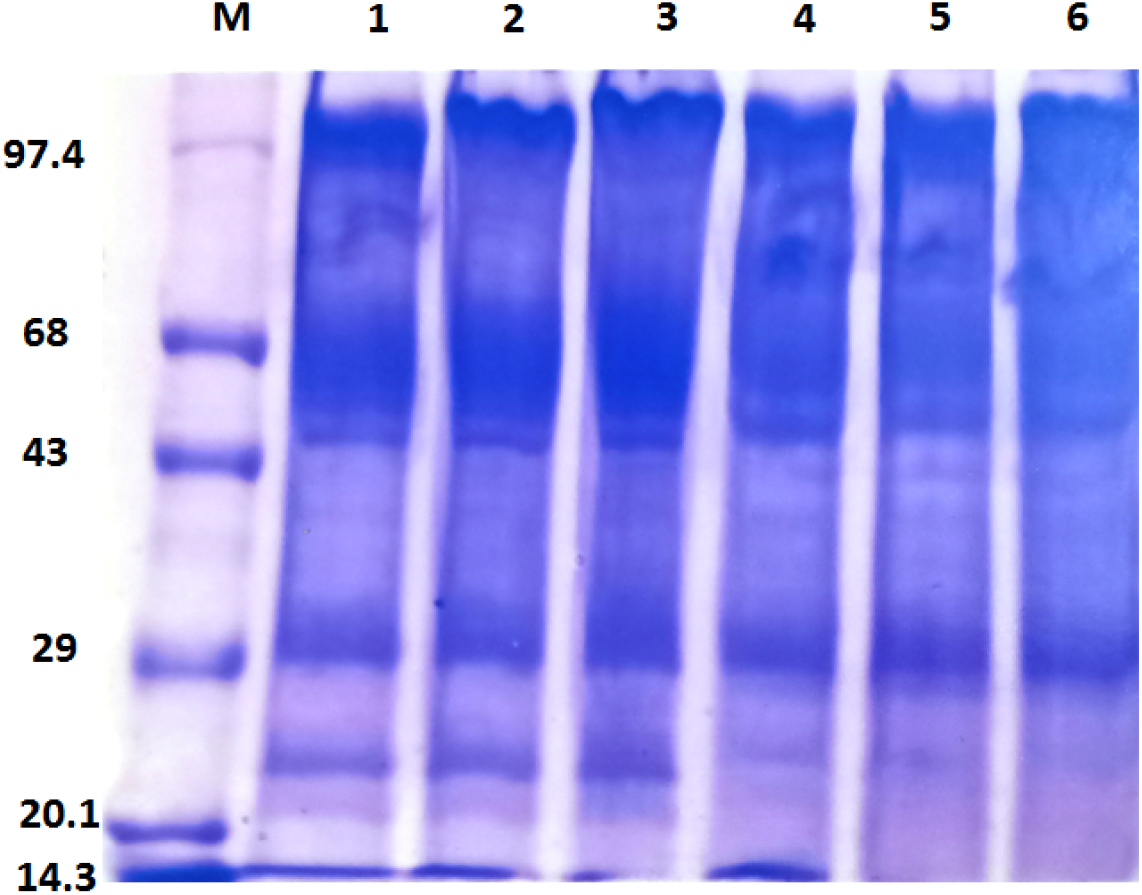
SDS PAGE of erythrocyte ghost membrane. Human and rabbit erythrocytes were prepared as described in the text and subjected to the SDS-PAGE. Lanes 1-3 contain human erythrocyte membrane 20, 25, and 30 ug protein respectively. Lane 4-6 contains rabbit erythrocyte membrane 20, 25, 30 ug protein respectively.

**Figure 6:**
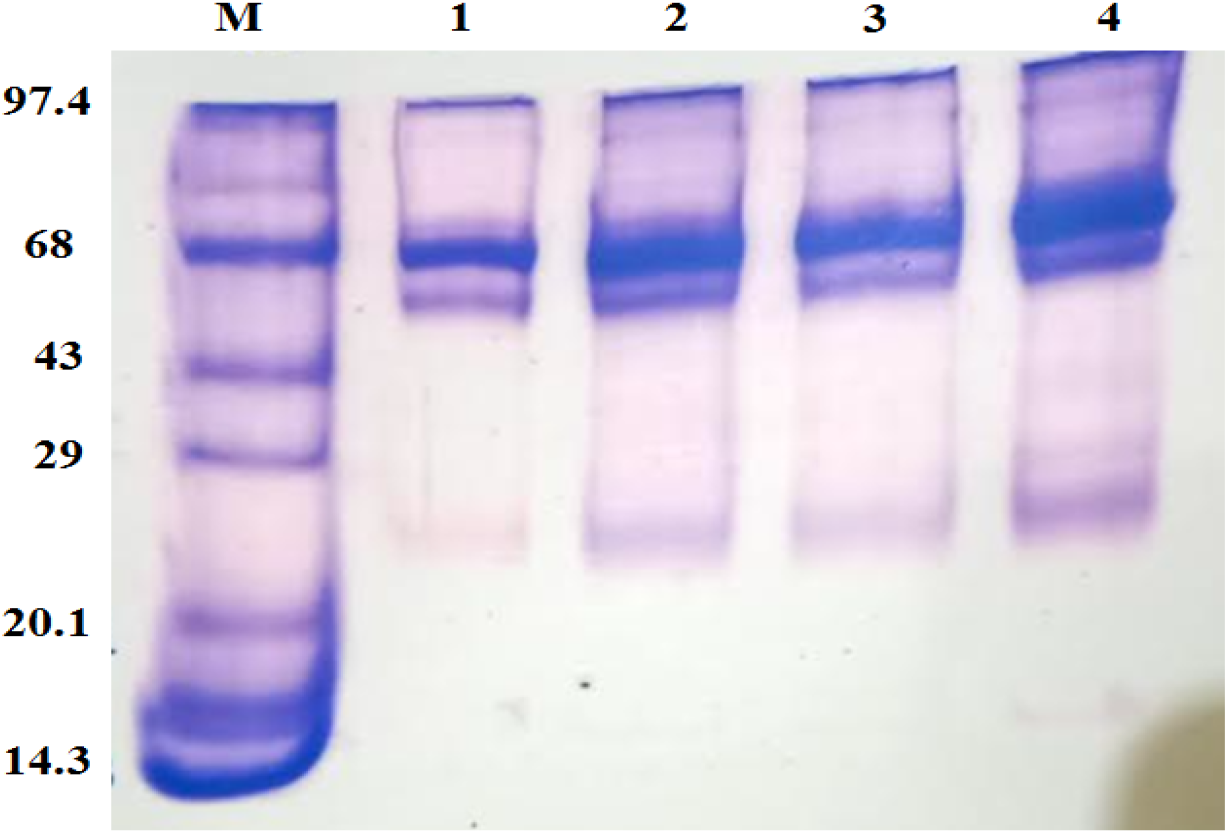
SDS PAGE of concentrated plasma by Sephadex-10. 0.25ml plasma was added 10 mg (lane 2), 20 mg (lane 3) and 50 mg (lane 4) of Sephadex-10. The samples were added and incubated for 2 hours centrifuges to remove Sephadex and the plasma subjected to SDS-PAGE 25 microlitres protein was applied in each lane. Lane l contains standard protein markers and lane 2 Plasma is not treated with Sephadex.

### 5.3 Immunizations with the invertase for Antibody response in the rabbits

The antibody reaction in rabbits after immunization with free invertase dissolved in PBS, entrapped in erythrocytes and loaded in erythrocytes and further entrapped in plasma beads was investigated. In all the rabbits receiving the antigen, IgG titres increased remarkably starting from day 12. Rabbits receiving the antigen formulation also showed almost similar profiles with values of day 20 being maximum and the titre going down by day 35. Interestingly, in the case of groups of rabbits receiving the enzyme entrapped in the erythrocytes and plasma beads, the IgG titre increased up to 28^th^ day and reduced just reasonably on 35^th^ day. This suggested the slow release of the enzyme from the beads. After 12 days of immunization, the rabbits’ antibody titre in receipt of erythrocyte entrapped invertase was maximum followed by those in the animals receiving plasma bead preparation and soluble enzyme in PBS (figure 7). After 20 days the antibody titre in animals receiving invertase dissolved in PBS declined while those in animals receiving erythrocyte encapsulated and plasma bead entrapped preparation rose further. Activity in the group receiving erythrocyte-entrapped enzyme still was maximum (figure 8). After 28 days the titre increase only in the group receiving the plasma bead preparation rose while those in other groups declined (figure 9). Although the titre fell in all groups after 35 days the decrease was minimum in the plasma bead group (figure 10).

**Figure 7:**
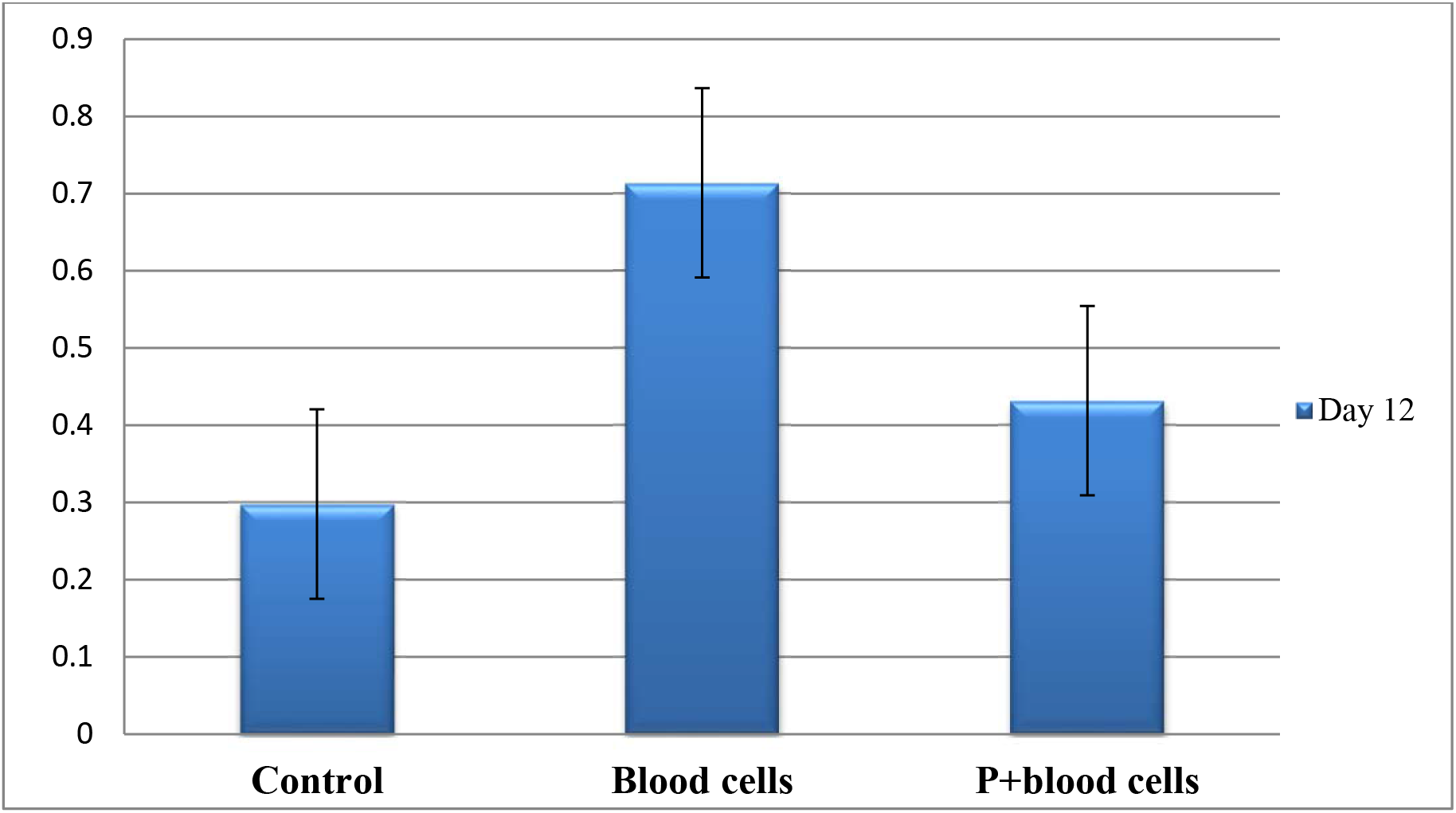
Antibody response in rabbits after immunization with invertase. Rabbit Groups were immunized with preparation invertase including at no cost enzyme dissolved into PBS; Invertase enzyme entrapped in the erythrocytes or invertase entrapped in erythrocytes and further included in plasma beads. Total IgG sera levels were calculated in the animals’ sera at the point of the period. All animals established 600 μgs of invertase like a single dose.

**Figure 8:**
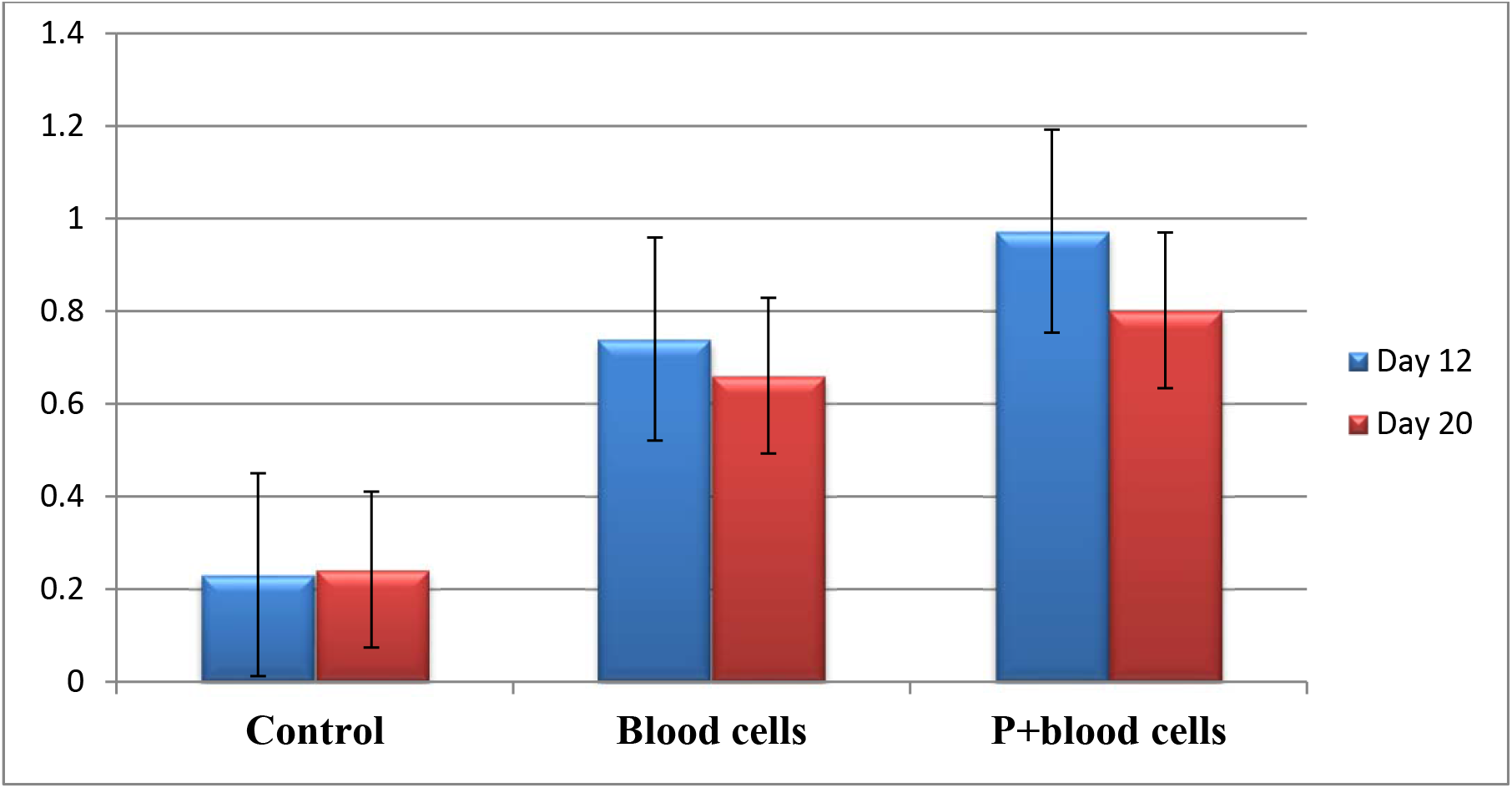
Antibody response in rabbits after immunization with invertase. Rabbit Groups were immunized with preparation invertase including at no cost enzyme dissolved into PBS; Invertase enzyme entrapped in the erythrocytes or invertase entrapped in erythrocytes and further included in plasma beads. Total IgG sera levels were calculated in the animals’ sera at the point of the period. All animals established 600 μgs of invertase like a single dose.

**Figure 9:**
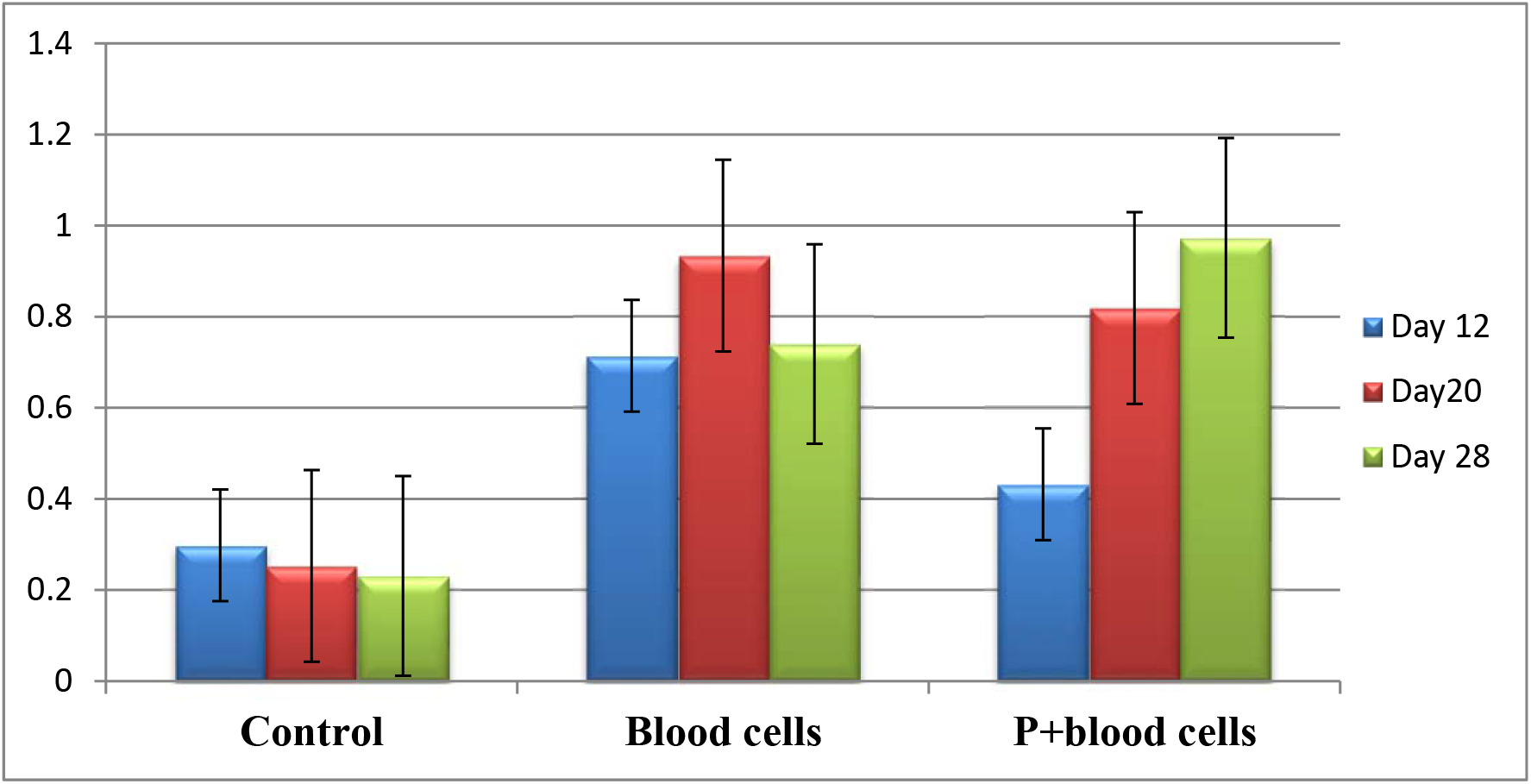
Antibody response in rabbits after immunization with invertase. Rabbit Groups were immunized with preparation invertase including at no cost enzyme dissolved into PBS; Invertase enzyme entrapped in the erythrocytes or invertase entrapped in erythrocytes and further included in plasma beads. Total IgG sera levels were calculated in the animals’ sera at the point of the period. All animals established 600 μgs of invertase like a single dose.

**Figure 10:**
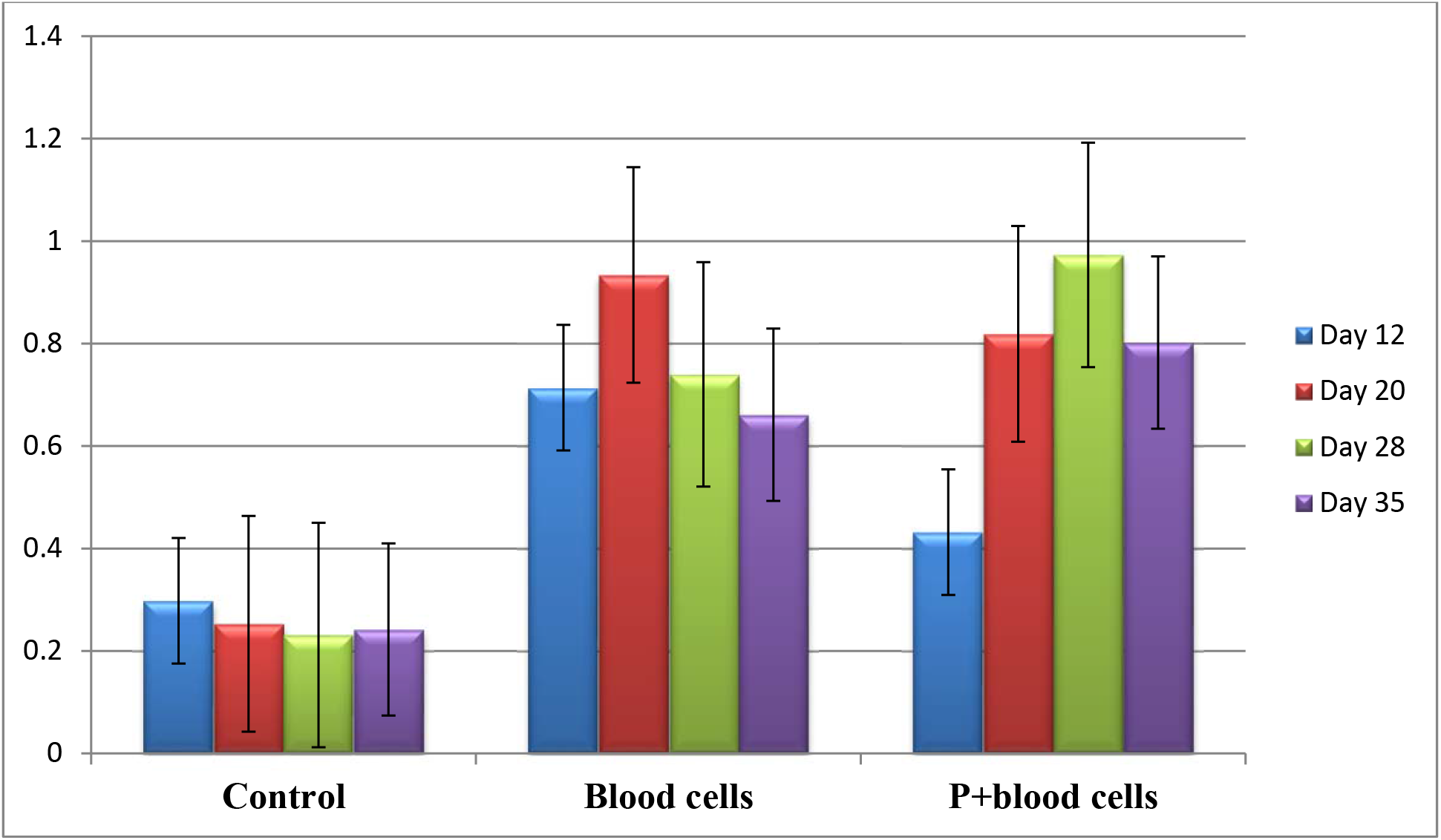
Antibody response in rabbits after immunization with invertase. Rabbit Groups were immunized with preparation invertase including at no cost enzyme dissolved into PBS; Invertase enzyme entrapped in the erythrocytes or invertase entrapped in erythrocytes and further included in plasma beads. Total IgG sera levels were calculated in the animals’ sera at the point of the period. All animals established 600 μgs of invertase like a single dose.

## Discussion

Vaccination is the most cost-effective strategy of controlling infections and has been and continued to be used for controlling and even eradicating infections. One of the challenges of successful immunization is the administration of multiple doses of vaccine. Also, most soluble antigens are not highly immunogenic unless they are administered along with the adjuvant only one adjuvant is used in case of human immunization. Biodegradable polymers have received remarkable alteration in recent years for the controlling release of antigens. The polymeric systems have been primarily employed to reduce the number of doses, to target the antigenpresenting cells, and out down to the quantity of antigen required. A number of polymeric matrices/microspheres have been needed for the entrapment of vaccine antigens (HIV, flu, HAV, allergens, etc). These include mainly synthetic polymers like poly (lactic/glycolic) acid and several derived from natural sources such as liposomes virosomes, etc. A number of reports are also available on fibrin polymers as drug delivery systems. A report on the preparation of antigens plasma beads without the requirement of additional thrombin or transglutaminase has been reported from this laboratory and this effort was assumed to examine the antigen delivery potential of the plasma beads in combination with erythrocytes bearing antigen. Reports are available on the usefulness of erythrocytes as carriers of drugs and antigens. Erythrocytes derived from rabbits, mice and humans could be loaded efficiently using the dialysis procedure. Although there was moderate variation in this amount of enzyme entrapped, an increase in this amount of increase added to the red cells resulted in increased entrapment. At a very high concentration of the added enzyme no further increase was observed in the amount entrapped (figure 2–4). The erythrocyte ghosts prepared from the cells were characteristic of all these bands of erythrocyte membrane (figure 5). All the proteins showed molecular weight reported in the literature suggesting a lack of degradation during the ghost preparation. Stable plasma beads were prepared using concentrated plasma protein and a high concentration of thrombin and transglutaminase. The results of immunization were interesting as shown in figures 7, 8, 9 and 10. The IgG levels of sera of rabbits receiving erythrocyte-entrapped enzyme were highest after day 12 followed by the group receiving the plasma bead entrapped erythrocyte encapsulated enzyme. Animals receiving the soluble enzyme in PBS contained the lowest amount of antibody. The level of the antibody rose in the groups receiving erythrocyte-entrapped enzyme till day 20 and declined gradually till day 35. Antibodies levels in the plasma beads administered group rose titre 28^th^ day and showed a small decline on the 35^th^ day. The antibody levels in the plasma beads administered group were markedly higher than the group that raised erythrocytes antigen. The antibody levels in the group receiving soluble invertase results were low throughout the investigation.

## CONCLUSION

These studies show that erythrocytes act as adjuvant and contribute towards the enhancement of the immunogens of the protein. Further, it is quite likely that plasma beads entrapment contributes to sustained delivery of the antigen and results in a remarkably high level of antibodies even after 35 days of immunization.

## AUTHOR CONTRIBUTIONS

Conceptualization, M.R.J.; Data curation, M.F.K., H.J.; Formal analysis, M.R.J,; Investigation, M.R.J.; Methodology, M.R.J.; Project administration, M.R.J.; Software, M.R.J., M.M., M.F.K., S. M., and H.J.; Validation, M.R.J.; Visualization, M.R.J.; Writing—original draft, M.KJ.; Writing—review and editing, M.R.J., M.M., M.F.K., S. M., and H.J.; All authors have read and agreed to the published version of the manuscript.

## ACKNOWLEDGMENTS

We would like to thank Prof. Saleemuddin (Retired), Interdisciplinary Unit of Biotechnology, Aligarh Muslim University, Aligarh, Supervisor (Master’s in Biotechnology) of M. Rizwan Jameel, for his generous help in this study.

## CONFLICTS OF INTEREST

The authors declare no competing interests.

